# Beyond Climatic Variation: Human Disturbances Alter the Effectiveness of a Protected Area to Reduce Fires in Tropical Peatlands of Sumatra, Indonesia

**DOI:** 10.1101/351817

**Authors:** Muhammad Ali Imron, Kirana Widyastuti, Ryan Adi Satria, Wiwid Prayoga, Subyantoro Tri Pradopo, Hatma Suryatmojo, Bertha Maya Sopha, Uta Berger

## Abstract

The occurrence of fires has frequently been used to highlight environmental hazards at regional and global scale, and as a proxy for the effectiveness of protected areas. In contrast, the mechanism behind wildfire dynamics in tropical peat land protected areas had been poorly addressed thus far. Our study provides a novel application of assessing fire patterns from a tropical peatland protected area and surrounding landscape. We investigated the importance of both climatic factors (top-down mechanism) and human interventions (bottom-up mechanism) on fire occurrences through analyzing 15-year (2001 - 2015) LANDSAT and MODIS images of the Padang Sugihan Wildlife Reserve (PSWR). Fire density along side road and canal construction were analyzed jointly together with the monthly and annual precipitation, and evidences of climatic anomalies. The reserve was effective in limiting fire occurrences from surrounding landscapes only in wet years. We revealed that peat fire patterns in the protected area and the landscape matrix emerged beyond climatic factors, and the distance from canal system could explain the fire occurrences. Our results show that it is essential to address processes at a landscape level, particularly at the surroundings of the reserve, in order to increase the effectiveness of fire protection, including the development of fire-prone classes maps.

## Introduction

Fire has a long historical relationship with humans [1]. Although less significant than other causal factors, fires are frequently used as an important indicator to evaluate the effectiveness management of protected areas [2, 3]. A common approach to manage fire in protected areas is applying active fire management or prescribed burning [4–7] which aims to reduce fuel availability for preventing and controlling wildfire [8]. Nowadays, a paradigm shift resulted in managers purposely burning grassland and forests to maintain the ecological mechanisms which drive ecosystem dynamics and diversity [7, 9].

Anthropogenic and natural factors lead to different patterns of fire occurrences in protected areas across various ecosystems. Fire density was found to be two times higher in non-protected areas than within protected areas in Myanmar [10] and Amazonian regions [2]. It has been shown that human intervention influences a protected areas susceptibility to disturbance. At a global scale, forest loss rates in protected areas is associated with high proportions of agricultural land in the country [11]. Managing anthropogenic factors which include fewer road construction, less human impact mechanisms [2] as well as fire-free land management [12] in the reserves has shown their effectiveness in reducing fire-driven deforestation. In contrast to fire occurrences in tropical areas, natural mechanisms caused higher fire density in protected areas than in non-protected areas in West and Central Africa [13]. More fires occurred in 59 percent of the area where deforestation rates dropped between 2000 and 2007, since more fuel was available for ignition [12]. Here, controlled ignition and active fire management are required to mitigate fuel availability.

Fire is a critical attribute to peatland but rarely occurred in the remote forest until the last 3000 years as anthropogenic factors started affecting peatland [14]. Peatlands in South-east Asia holds 26 million ha which represents 69% of Tropical Peatlands globally [15–17]. In Indonesia, peatlands are vital ecosystems, covering 18-20 million hectares, or 10% of terrestrial area which significantly contribute to primary sources of wood and livelihood of local people [18].

In the last two decades, Indonesia has been experiencing a drastic shift in peat-land dynamics, from being frequently, from being frequently inundated and moist-ecosystem into human-made drained-ecosystem. This ecosystem change brings consequences for the frequent occurrences of peat-fires and subsequent health, environmental and biodiversity problems [17]. The recent fires of 2015 in Indonesia were the highest since the 1997 megafires and almost the whole Sumatra island was engulfed by smoke, while South Sumatra province holds the second highest number of hotspots amongst the Indonesian provinces during 2015 [19]. Particularly in Sumatra island, the combination of deforestation from adjacent areas [20], the rapid expansion of palm oil plantation, and the use of fire for land preparation [21] significantly increase threats in protected areas as well as biodiversity conservation of selected species [22]. Studies on the effectiveness of protected areas in a peatland to reduce wildfire are still limited. Fire in Indonesian peatland has been intensively studied through remote sensing data to predict fire effects [23–25], peat hydrology [5], fire database management [26], effect of fire on bio-physical attributes attributes [24, 27, 28], and peat restoration planning [29]. Nevertheless, studies with emphasis on protected areas are still rare.

Fire management in Indonesia has different characteristics from the majority paradigm. While the use of fire is encouraged in savannas [7, 9], efforts on fire prevention are still relevant to address ecological issues in Indonesia, particularly in peatlands. In both cases, understanding causal mechanism of fire occurrence is important. The patterns and causal factors of fire in a protected area might help reveal more information to guide effective fire management. In addition, current studies often overlook the importance of peatland protected areas on reducing fires in the landscape.

We aim to gain insight into pattern and causal factors of fires through the comparison of fire occurrences within and surrounding the Padang Sugihan Wildlife Reserve (PSWR), a protected area in South Sumatra which is dedicated for biodiversity conservation. We observed a 15-year period of fire occurrences data to provide a basis for evaluating the effectiveness of peatland protected area to reduce fire intensity from the surrounding landscape.

**Fig. 1.**
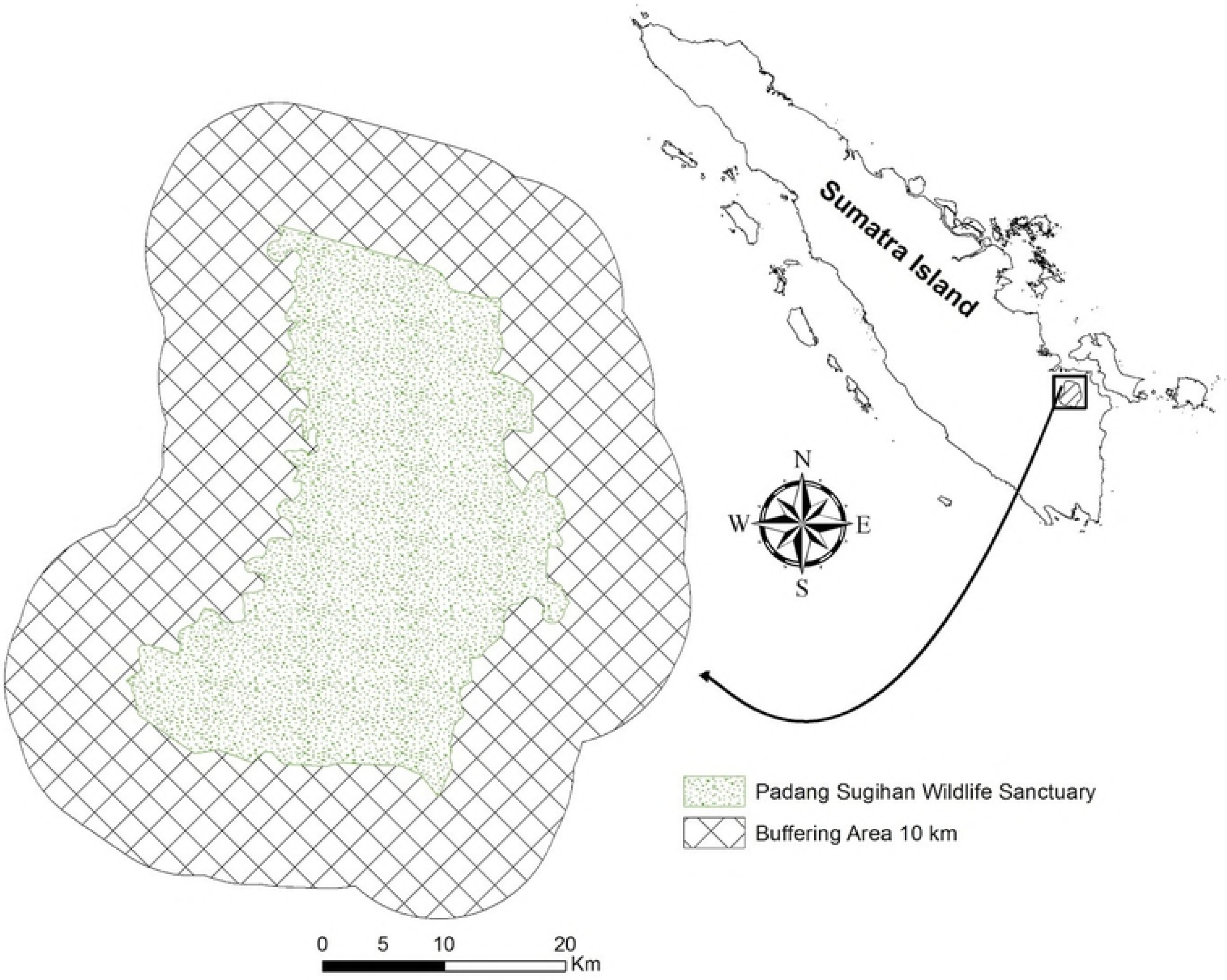
Padang Sugihan Willdife Reserve and its surrounding area (10 km buffer) in South Sumatra Province of Indonesia.

## Materials and methods

### Study Area

Padang Sugihan Wildlife Reserve (PSWR) is among protected areas managed by South Sumatra Balai Konservasi Sumber Daya Alam (BKSDA)/ Natural Resources Conservation Agency, Ministry of Environment and Forestry, Republic of Indonesia. This reserve aims to protect the remaining Sumatran elephant (*Elephas maximus sumatranus*) populations in this province. PSWR occupies an area of 881.48 km^2^ which is dominated by peatland. Sugihan river lies to the west of the reserve, while Padang river serves as the border to the east. To the south of the reserve, lies Butung river, and a canal is built to the north (Fig 1). Administrative boundary maps of South Sumatra (http://www.bakosurtanal.go.id/peta-rupabumi/) and Padang Sugihan Wildlife Reserve maps were used in this study and obtained from BKSDA.

### Fire detection and Environmental Studies

Fire detection was processed using Active Fire data provided by the Moderate Resolution Imaging Spectrometer (MODIS). It provides the geographical position of fires within 24 hours of monitoring [13, 30] as well as 1-2 day return interval which is useful to monitor the change of fire detection at any given location [30]. We selected MODIS active fire data in the period between 2001 and 2015 with a confidence level of 80%. Since Indonesia is influenced by El Nino / Southern Oscillation (ENSO), we categorized the fire data to wet and dry periods according to NOAA Climate Prediction Centre [31].

To compare fire occurrences within and surrounding the reserve, we calculated fire frequency within the reserve and within 10 km surrounding its border. A buffer zone of 10 km was chosen because it is equal to the longest distance from the border to the reserve center point (Fig 1) and is commonly used to assess protected area effectiveness [32]. The buffer size was 1,781.68 km^2^, combined with PSWR (881.48 km^2^) it occupies a total area of 2,663.16 km^2^. We divided our study site into 1 × 1 km^2^ grid cells to identify whether fires can occur in the same area. Recurrent incidences of fire in certain cells over the observed period are classified as repeated fire.

We plotted fire occurrences and daily precipitation to extract relational patterns between the two factors. To understand the impact of reserve border in reducing fire density, we compared fire frequency within the reserve and 1 km surrounding it. We used the buffer function from ArcGIS 10 to measure fire density in each distance interval. The distribution of fire for the period of 2001 - 2015 was calculated with Kernel Density Estimation using ArcGIS 10.

To understand the relation between fire occurrence and precipitation, we obtained rainfall data during 2001 - 2015 from the nearest weather station, Sultan Badaruddin II Airport in Palembang. Anthropogenic factors were derived from roads and canal system within the reserve and surrounding areas during the period of 2001 - 2015. We manually digitized satellite images from Landsat 7 and Landsat 8 (https://landsat.usgs.gov/) to detect canal and road networks for the same year. To test the relationship between monthly precipitation and number of rainy days on the fire occurrences, we performed a Chi-square tests.

To test the roles of various environmental factors on the probability of fire occurrence, we used Normalized Density Vegetation Index (NDVI), Enhanced Vegetation Index (EVI), distance from canal (dist-canal), and distance from road (dist-road) for the calculation of binomial generalized linear model with R statistics version 3.0.0. We used the model to develop a fire-prone probability map for both dry and wet years to provide an early indication for the management of the reserve using Arc GIS 10.

**Fig. 2.**
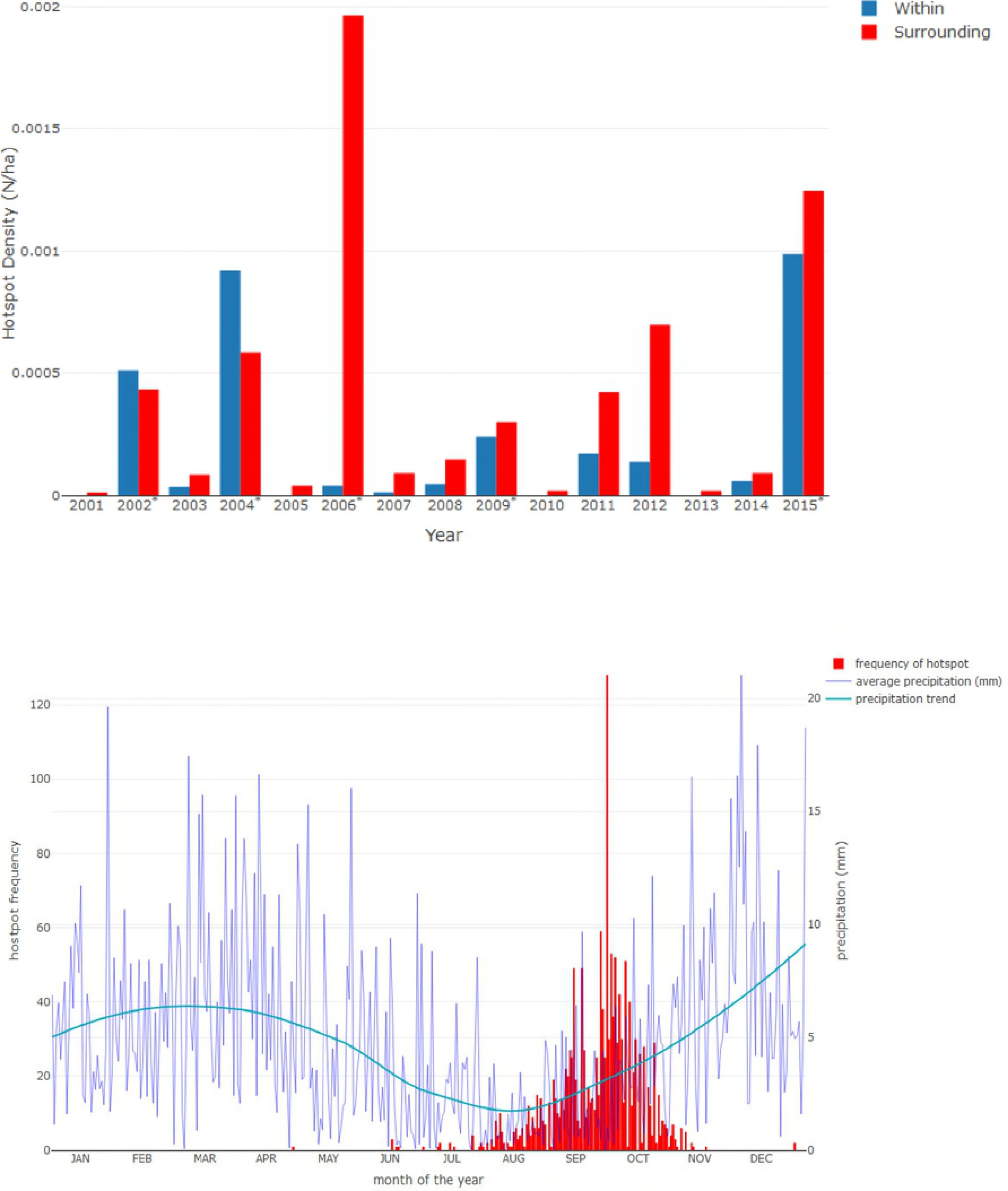
Fire densities within and surrounding Padang Sugihan Wildlife Reserve during 2001 to 2015. Fires within the reserve tend to be lower than those within the surrounding. However, anomalies occurred during 2002 and 2004 during dry periods (above).The temporal relationship between the average daily fires and the average precipitation during 2001 - 2015 (below).

**Table 1.** Relationship between average monthly precipitation and number of rainy days each month, with number of hotspot during 2001 to 2015 in Padang Sugihan Wildlife Reserve.

| Variable | $\tilde{\chi}^2$ | p-value |
| --- | --- | --- |
| **All year (n = 180, df = 2)** |  |  |
| monthly average precipitation | 0.07 | 0.0004337 |
| monthly number of rainy days | 0.13 | 6.40E-07 |
| **Dry periods (n = 60, df = 2)** |  |  |
| monthly average precipitation | 0.13 | 0.00426 |
| monthly number of rainy days | 0.31 | 3.52E-06 |
| **Wet periods (n = 120, df = 2)** |  |  |
| monthly average precipitation | 0.04 | 0.02166 |
| monthly number of rainy days | 0.06 | 0.008715 |

## Results

### Temporal and Spatial Patterns

Fire densities within the reserve and the buffer zones can be classified into two pronounced periods which are 2001 - 2006 and 2007 - 2015, both characterized by a steady increase of observed fires per hectare (Fig 2) with the highest density recorded in 2006. Fire densities were lower within the PSWR than in the buffer zone except in the two dry years (2002 and 2004).

The role of precipitation dynamics on fire emergence was clearly shown with a functional delay of monthly precipitation decrease in July followed by increasing numbers of wildfires from August to September (Fig 2). Additionally, the average monthly precipitation and number of rainy days shows associations with number of hotspots during dry years, wet years as well as for all years during 2001-2015 (Table 1).

The fire occurrence in PSWR and its surrounding area was most frequent during the dry season from July to November. Only a few anomalies of fire occurrence were observed during May and December. Fires (n = 180) were able to emerge when no rain occurred or up to 37 consecutive dry days happened prior to the ignition (median = 8 days). Whereas, the duration of fire (n=180) to maintain burning was between 1-14 days with a median of 1 day.

During the observed period, most fire incidences, within and surrounding PSWR, occurred only once. We found fewer grid cells in which fire occured twice or more, and only few were burnt more than seven times. It was clearly shown that fire frequencies was significantly lower within the reserve. In the 15 year period, a very small portion from the surrounding of PSWR was burnt annually while within the reserve, only 4 repeated fire occurred at most (Fig 3).

**Fig. 3.**
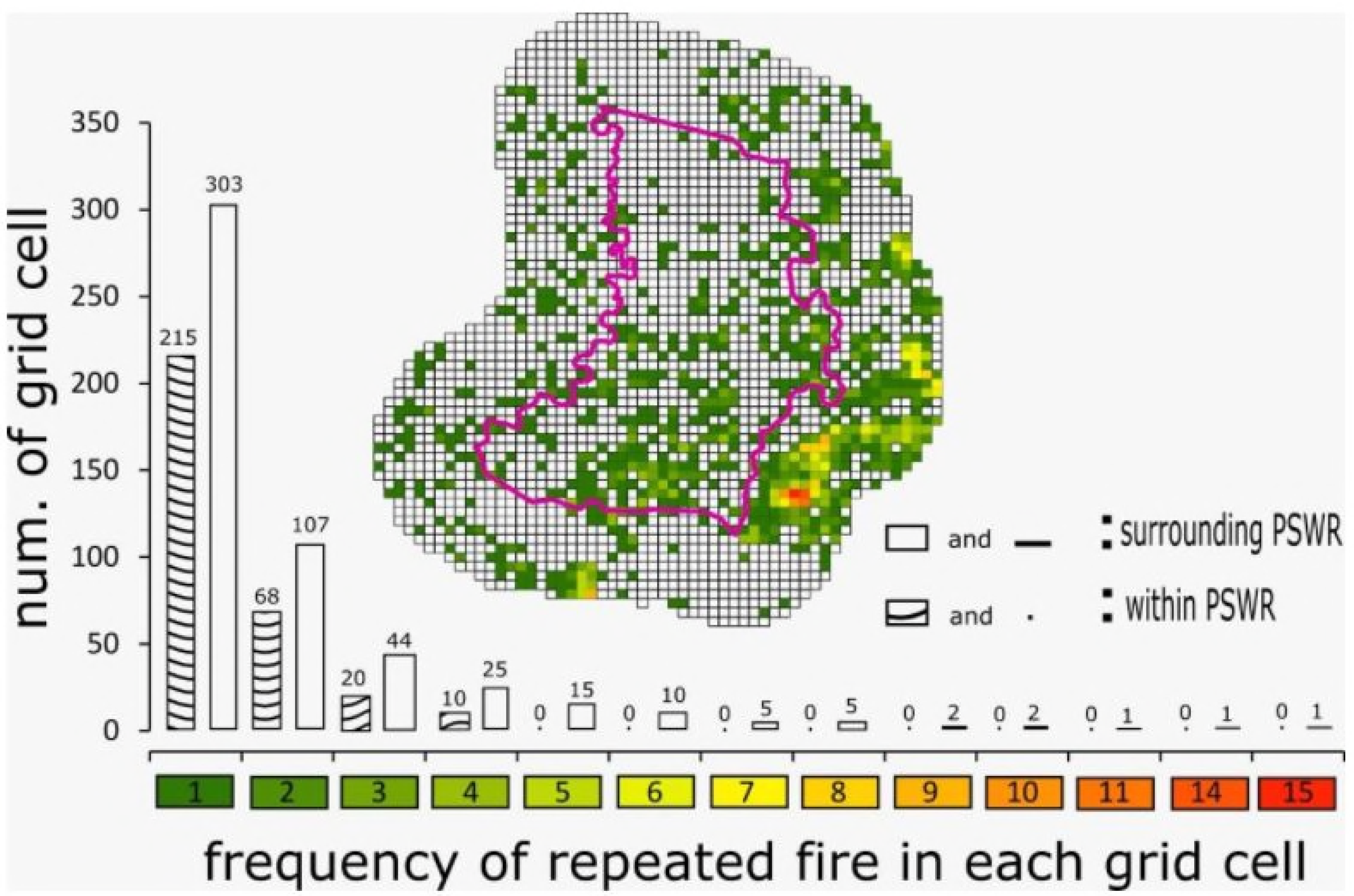
Number of grid cells in each frequency of repeated fire within the Padang Sugihan Reserve and surrounding between 2001 and 2015.

**Fig. 4.**
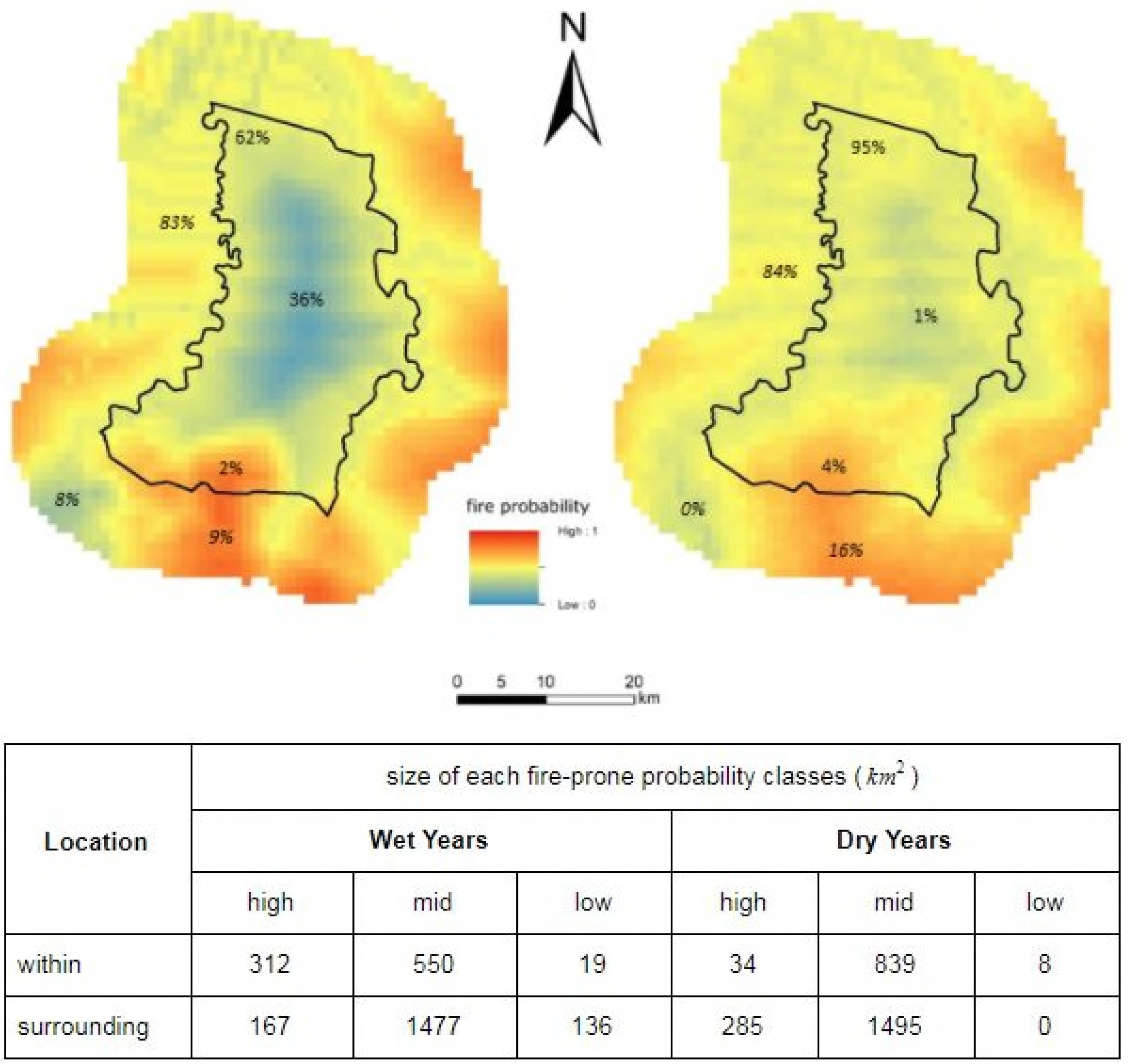
Hotspot density during wet and dry years within and surrounding the reserve border (above). The frequency of hotspot as function of distance from canals and roads (below)

In the wet years, fire density was higher surrounding PSWR, whereas in dry years fire occurs proportionally within and surrounding the reserve. Fire density decreased from the reserve border to the core during the wet years, whereas an unclear pattern was shown in dry years. During wet years the density of fire within 1 km from the reserve border was low, and at a gradually further distance fire occurrences were found to be increasingly rare. In addition, fire density outside the reserve tended to increase with further distance up to 6 km from the border (Fig 4). Conversely, in dry years we could not see the gradual reduction of fire occurrences from the border to both within and surrounding the reserve. In dry years, the irregular pattern of fire density increment within PSWR was similar to which surrounding the reserve. However, a relatively lower density of fires was identified near the reserve border.

During dry years, distance to canals, distance to roads, NDVI, and EVI have a profound effect on fire presence probability, whereas in wet years the most critical factor affecting fire occurrence is the distance to roads and distance to the canal (Table 2). During wet years fires rarely occurred within the protected area, except for a small portion of 2003, 2008, 2011, 2012, and 2014. During warm or dry years, the fire spread throughout the landscape including within the reserve. We highlighted that human disturbance as represented by canal and road systems have a pronounced role on fire occurrences during dry and wet years. A reverse J pattern was shown, indicating a decrease of fire incidents as the distance from canals and roads increase. Meanwhile, the distance from reserve border has a less clear impact on fire frequency (Fig 4).

Our binomial generalized linear model clearly shows a sudden shift of reduced size of low fire-prone probability class within the PSWR from wet years (36%) to dry years (1%). In contrast, a sharp inclination was shown for the middle fire-prone probability class from wet years (62%) to dry years (95%) within the PSWR (Fig 5). The proportion of each class was not changed significantly for the surrounding PSWR from wet to dry years. However, during the dry season, none of the surrounding areas of PSWR had the low fire-prone probability class. Our findings highlight that surrounding of the PSWR is continuously under the threat of fire, while within PSWR can protect some areas during wet years but not during dry years.

## Discussion

Various studies on the effectiveness of protected areas to mitigate fires exist, and provide mostly large-scale spatial analysis [2, 3, 10, 33]. Our approach to analyzing peat-fire occurrences at a local scale offers new insight into the detailed temporal and spatial patterns of fires within and surrounding a reserve and serves as the first local-specific case for a peatland protected area in Sumatra. This study demonstrates that climatic variation is not necessarily sufficient to explain fire emergence within and surrounding a protected area. We have demonstrated that both top-down (seasonal variation) and bottom-up (human disturbance) mechanisms play crucial roles on patterns of peat fires occurrences, and thus affect the reserve effectiveness in reducing fires. We discuss our key findings and conclude with a commentary on implications for protected areas’ management and biodiversity conservation.

**Fig. 5.**
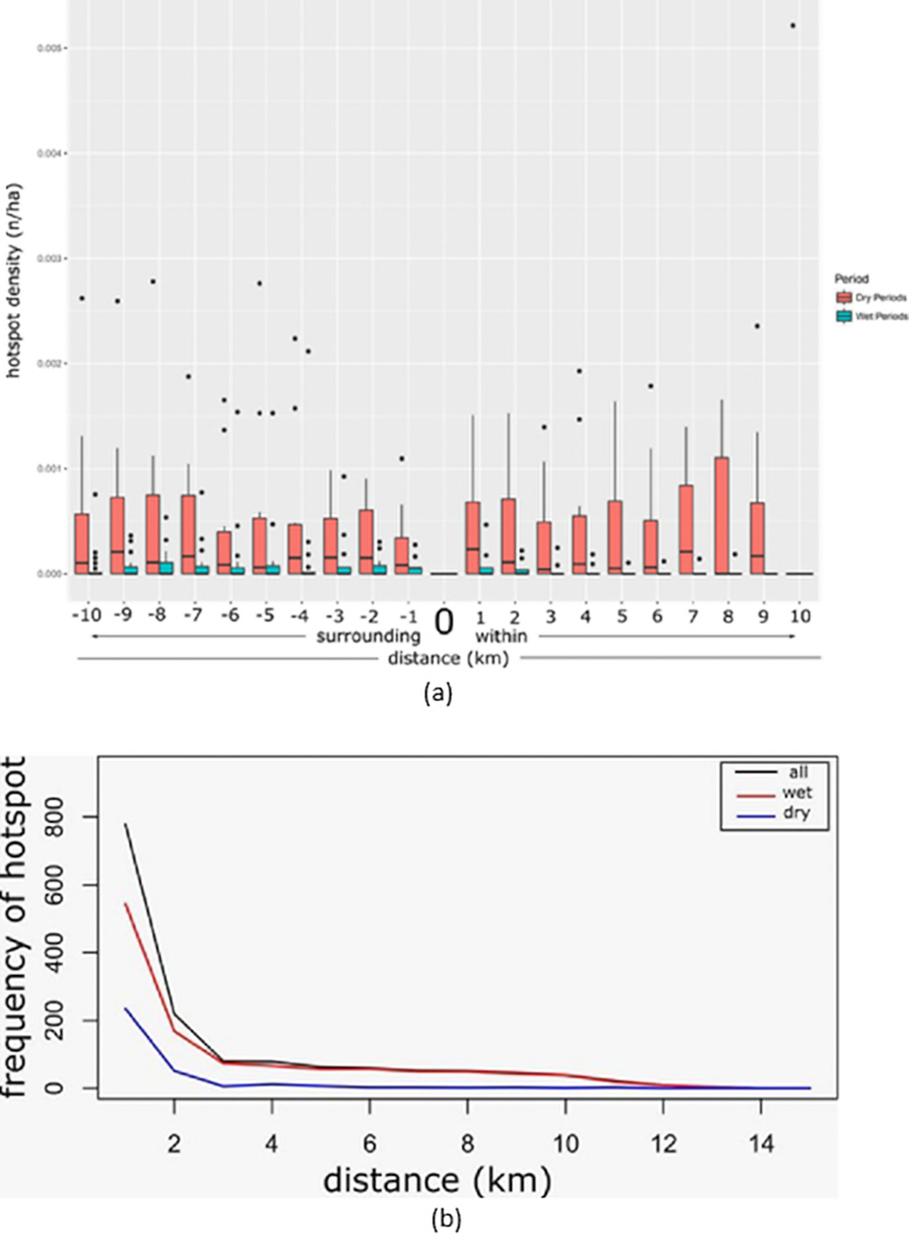
Predicted map of fire-prone probability class and its proportion for wet years (left) and dry years (right) from the binomial generalized linear model. The probability of fire occurrence is represented by different colors ranging from red (high) to blue (low). The area size of each fire-prone probability classes (below) compared in both wet and dry years.

**Table 2.** Environmental co-variates for the probability of fire presence in Padang Sugihan Wildlife Reserve for dry and wet periods from binomial generalized linear model

|  | Estimate | Std. Error | z-value | Pr(>\|z\|) | Sign. Code |
| --- | --- | --- | --- | --- | --- |
| Dry periods |  |  |  |  |  |
| (intercept) | 3.16E-01 | 4.37E-01 | 0.722 | 4.70E-01 |  |
| dist_roads | -3.41E-05 | 1.40E-05 | -2.442 | 1.46E-02 | * |
| NDVI | -2.80E+00 | 1.05E+00 | -2.663 | 7.74E-03 | ** |
| EVI | 3.09E+00 | 1.15E+00 | 2.675 | 0.00748 | ** |
| dist_canal | 9.836E-05 | 1.598E-05 | 6.156 | 7.46E-10 | *** |
| Variable Importance |  |  |  |  |  |
| dist_canal : 6.156 |  |  |  |  |  |
| evi : 2.675 |  |  |  |  |  |
| ndvi : 2.663 |  |  |  |  |  |
| dist_roads : 2.442 |  |  |  |  |  |
| Wet periods |  |  |  |  |  |
| (intercept) | -2.02E-01 | 1.37E-01 | -1.477 | 1.40E-01 |  |
| dist_roads | -2.11E-04 | 4.23E-05 | -4.991 | 6.01E-07 | *** |
| dist_canal | 1.36E-04 | 2.56E-05 | 5.334 | 9.62E-08 | *** |
| Variable Importance |  |  |  |  |  |
| dist_canal : 5.334 |  |  |  |  |  |
| dist_roads : 4.991 |  |  |  |  |  |

Climatic anomalies such as El Nino and the Indian Ocean Dipole pattern contribute significantly to fire incidences in Indonesia [21]. We have demonstrated top down perspectives act as a casual causative explanation of fire occurrences. We observed the impact of seasonal variation of fire vulnerability by comparing fire density in wet and dry years. Our analysis revealed a clear indication of different fire regimes during dry and wet years, which contributes substantially in fire-prone area predictive map development. Therefore, early-warning map development systems should consider the precipitation dynamics in both wet and dry years (Fig 5). During wet years with more precipitation and rainy days, the reserve area was able to limit fire. However, in dry years, when the rainfall is low the presence of protected area borders as a mean of land-use management was ineffective to prevent burning events [34], causing an increase of fire occurrences within and surrounding the reserve.

Our findings confirm that the accumulated rainfall and the length of the dry season influence the annual area burned in protected areas [35]. we provided here fine-scale fire distribution information in the landscape which is essential to understand the causal factors of fire in the reserve and surrounding landscape. From the bottom-up perspective, we analyzed physical and biotic factors which which are of vital for fire occurrences. Our results show that the distance to canal and distance to roads are important factors in the emergence of fire, both in wet and dry years. The construction of canals and roads represent human intervention, which is a primary trigger in fire ignition. In addition, the presence of canals implies a higher probability of drought in the surrounding area; which affects its vulnerability to fire ignition and spreading [36–38].

Vegetation index derived from satellite images have been commonly used for measuring fuel availability, which can also be used as a vegetation indicator for its health and susceptibility to fire [39]. Our study used Normalized Difference Vegetation Index (NDVI) and Enhanced Vegetation Index (EVI) on an annual basis. During dry years soil moisture content is decreased, thus affecting vegetation health; NDVI and EVI values are very low which are additional factors affecting the vulnerability of the peat area to fire. Yet our study has not explored whether any seasonal changes of NDVI influence the susceptibility to fire due to limited amount of clear satellite images (i.e less than 10% cloud cover) obtained per-season. We concluded that due to the lack of multitemporal NDVI series, we could not ascertain vegetation phenology. Hence, the complex interactions between vegetation, land use and climate characteristics cannot be explained [40].

In our study, the approach to use NDVI and EVI as vegetation indexes is limited by the relationship between fire; these indexes cannot be understood as a cause and effect relationship. We collected environmental variables (including NDVI and EVI) in the same year of the fire without considering the temporal relationship of fire occurrence. However, environmental variables would be better observed prior to a burning event to aid in determining cause-effect relationships between vegetation indexes and fire occurrences. Vegetation seasonal variability which indicates the shift from moist to dry conditions can help to identify the primary bioclimatic drivers related to fuel dynamics [40]. Implementing high-quality resolution of the multi-spatiotemporal image to depict the role of vegetation biomass will provide a substantial contribution to understanding the spatial pattern of fire [41].

The fire occurrences follow a functional delay response towards precipitation during 2001-2015 in the PSWR. The high fire occurrence during wet years in 2005 and 2010 indicates that natural drought processes could not sufficiently explain fires in the reserve. Since fires still occurred during the wet season, it is suspected that other factors may explain the temporal pattern observed in this research. The functional delays of fire occurrences after days without rain shows a combination of the bottom-up and top-down mechanisms. Since fire in peatlands are mostly anthropogenic, the functional delay represents the role of humans in igniting fires. We were not able to simulate both top and below-ground fire spreading mechanisms, further exploration of farmers decision-making process to burn land for agricultural conversion is required, as fire is employed as a tool in agricultural-conflict scenarios pertaining to land rights/ownership [42–44].

We rarely found repeated fire in an area during the 15-year period and very few recurrent fires in the same grid cell. This phenomenon implies that the peatlands self-restored within the 15 year period and intermittent fire was avoided. It is commonly believed that isochronal fire is a function of land conversion into agricultural uses [38]. The absence of recurrent fires means that degradation of the PSWR only occurred recently. Commonly fire use by locals only occurs when initially opening/preparing land during the onset of the planting period [41–44]. When the initial fires occurred anthropogenic activities were also present which effect flooding rates in the grid cells; therefore cessation of land conversion prevented recurrent fires from being ignited. [45]. Extension of observations, to exceed a plantation rotation may provide better understanding of fire use for land conversion and resulting recurrent fire.

Protected area ineffectiveness also stems from the presence of canal networks, whilst such networks provide access they also drain peatlands and increase the likelihood of it burning [36–38]. This evidence shows the bottom-up mechanism. Canal construction as a part of peatland conversion is commonly followed by the construction of drainages, and serves as a proxy of human disturbance to predict fire in the PSWR and surrounding landscape. The patterns of the fire illustrated the canal scars still play essential roles on fire during dry years as waterways give access to fire, representing the human-induced activities [46, 47] including forest conversion into plantations [48]. Furthermore, canals and roads as a proxy of human presence serve as valuable predictors of fire occurrence in our study area, as well as elsewhere [2, 49]. Our approach, however requires additional land cover and land use analysis [50–52] to assess causal factors of fire in the surrounding landscape of PSWR. Recent perspectives pertaining to the role of hydrological drought on fire concurrences indicate additional studies on such dynamics in disturbed and undisturbed areas may influence our findings. Additionally research should seek to ascertain the motivations of anthropogenic land burning [53] to aid in understanding humans role in dry year fire amplification [2, 46, 54].

Fire presence within the reserve during wet and mostly dry years indicate the reserves biodiversity is under serious threat through habitat destruction and isolation within their boundaries [49, 55]. Responses of biodiversity, such as mammals, to and following fire incidences provide insight of the reserves effectiveness in maintaining such biodiversity, which was unfortunately beyond the scope of the present study. A particular emphasis on the behavioral response of Sumatran elephants as a flagship species in PSWR, using GPS collaring [56]and population projection studies [57] would gain insight into the effect of fire on endangered species conservation. Furthermore, population viability analysis for the elephant in the reserve and surrounding area using a modeling approach [22, 52, 58] will provide guidance for management alternatives, currently lacking such data.

This study provided insights on spatial and temporal fire occurrence predictors, including human interference and climatic variations. Nevertheless, knowledge on the detailed mechanism of fire occurrence and its causal factors remain incomplete. Individual/Agent-based modeling is one promising approach to understanding the complex phenomenon of wildfire, as well as predicting fire-prone areas in the future [59–63]. Development of such models would be beneficial for applied science development on fire ecology and management. Predictions from such dynamic models, incorporating bottom-up and top-down processes would allow these complex processes to be incorporated, providing dynamic information applicable to management fire mitigation policy development. Also since vegetation and social conditions are highly dynamic in the tropical peatlands [42], using such dynamics model with relatively short temporal scale will enhance the prediction.

### Implication for Protected Areas Management

Protected areas in Sumatra have shown their effectiveness lowering the deforestation rates in comparison to adjacent unprotected areas during years of 1990 to 2000 [20]. However, our study shows that a protected area in a peatland ecosystem was not devoid of dry season fires. We also noticed the surrounding landscape becomes critical for the protected areas if adjacent areas could not hold fire occurrences. The surrounding area is key in mitigating fire, particularly during dry years. In addition, raising awareness of people surrounding the reserve to use fires wisely and practice sustainable peatland management, and supporting the works of Masyarakat Peduli Api (Fire-Concerned Community) in each village surrounding the reserve should be improved through collaboration with local government and private sectors. This will be crucial to the effectiveness of the PSWR to mitigate fire incidences.

The PSWR has crucial roles on biodiversity conservation in Indonesia as habitat for the endangered species of Sumatran Elephant. This subspecies of elephant has a large homerange [57] which consequently needs various habitat types and larger habitat size. The landscape approach of considering composition and quality of land surrounding a protected area has emerged recently in biodiversity conservation [22], and urgent implementation has been recommended. Encouragement of privately protected areas [64, 65] and village-based protected areas such as a sacred forest [66, 67] in the surrounding area of the reserve may enhance the effectiveness of biodiversity conservation and fire mitigation. In case of fire incidents within the PSWR, management should ensure safe areas from fire as a wildlife refugia system are present [68, 69].

Since the PSWR is also surrounded by canal systems and rivers for local transportation access to the reserve is relatively easy. Suggestions or the proposal to change the reserve into mixed-used protected areas to deal with pressure from agricultural and timber extraction [70] should be carefully considered. Attention to evaluate the effectiveness of the BKSDA Sumatera Selatan to manage the reserve [71–74] with particular emphasis on fire management;should be considered; prior to contemplating changing the status of protected areas into other protection types or other land use types. Strengthening the ability of rangers to detect and handle fires when they occur, will also enhance the effectiveness of protected areas’ management. Simultaneusly peatland restoration, mainly from the scars of canals within PSWR, using physicals approaches such as re-wetting through canal blocking and other methods [75–78] will help reduce the fire incidences.

## Acknowledgments

This project is funded by KEMENRISTEKDIKTI 2254/UN1-P.III/DIT-LIT/LT/2017. BKSDA Sumatera Selatan provided entry permit to the Padang Sugihan Reserve. The authors are thankful to Bukhori Akhmad for supporting the field works and Matthew Gardiner for proofreading the manuscript.

